# Use of dye sensitizers for increased photoacoustic mechanosensation

**DOI:** 10.1101/2023.11.15.567262

**Authors:** Tarek Rafeedi, Laura Becerra, Nicholas Root, Baiyan Qi, Lei Fu, Guillermo Esparza, Yi Qie, Lekshmi Sasi, Romke Rouw, Jesse Jokerst, Darren Lipomi

**Affiliations:** Department of Nano and Chemical Engineering, University of California San Diego, 9500 Gilman Dr., La Jolla, 92093, CA, USA; Department of Electrical and Computer Engineering, University of California San Diego, 9500 Gilman Drive, La Jolla, 92093, CA, USA; Department of Psychology, University of Amsterdam, Nieuwe Achtergracht 129-B, WT, 1018, Amsterdam, Netherlands

## Abstract

The photoacoustic effect refers to the generation of pressure waves in matter stimulated by light[1]. In the context of radiology (i.e., photoacoustic imaging) waves generated by pulsed laser light are detected by an ultrasound transducer[2–4]. It has been shown that photoacoustic waves produce a mechanical, tactile sensation in humans on bare skin[5]. In a series of psychophysical experiments, performed with both medical grade and off-the-shelf pulsed light systems, participants could detect, categorically describe, and discern the direction of travel of pulsed optical stimuli with the use of a dye as an optical absorber on the skin. To a large extent, the sensations were perceived as localized vibration on the glabrous surface of the fingers, when sensitized with the thin film of dye. This form of sensory stimulation demonstrates an enhanced non-contact, non-optogenetic, in situ activation of the mechanosensory system. This modality of sensation may provide a tool that leads to new insights in psychology, neuroscience, mechanobiology, and the health sciences. Finally, it has many advantageous characteristics for human interaction with artificial environments, as optical signals can be projected onto the skin across distances.

---

Mechanical forces at the surface of the skin are transduced to the brain by the cutaneous end organs and afferent fibers of the somatosensory system [6, 7]. Along this pathway, signals are encoded and ultimately categorized as familiar conscious percepts, e.g., pressure, vibration, stretch, and pain. In natural environments, mechanical forces great enough to initiate action potentials must be exerted by a solid or fluid acting directly on the surface of the skin.

The photoacoustic effect, discovered by Alexander Graham Bell in the 1880s, involves the use of pulsed light to generate pressure waves in matter[1]. The effect has been used for decades in materials research to measure dissipation and storage mechanisms in solids[8–10]. More recent applications have used the effect in concert with high-frequency ultrasound to create images and 3D maps of tissues using endogenous absorbers, e.g., hemoglobin, or exogenous absorbers [2–4], as sensitizers. The penetration depth of red or near-IR light, along with the lack of scattering experienced by acoustic waves in biological tissue, has made photoacoustic tomography a robust tool for imaging with high resolution at depths of centimeters[11]. In the last decade, it has been demonstrated that pulsed laser light can evoke tactile sensations on the skin mediated by thermoelastic effects[5, 12]. Incident pulsed light is converted to thermal energy, inducing thermal expansion and subsequent relaxation forming stress (acoustic) waves.

The central hypothesis of this work is that the stress waves generated in the skin by the photoacoustic effect will produce a more robust and enhanced tactile sensation with a thin layer of an optical absorber (dye) on the skin, as shown in (Fig. 1a,b). That is, mechanical signals with frequencies below approximately 1000 Hz can in principle be detected by the mechanosensory neurons of the human skin. For example, the Pacinian corpuscle is most sensitive around 250 Hz extending to 1 kHz[6]. The Meissner’s corpuscle, responsible in part for sensations of fine texture, is most sensitive in the tens of Hz, and the Merkel cell-neurite complex is sensitive at the level of a few Hz [13, 14].

**Fig. 1.**
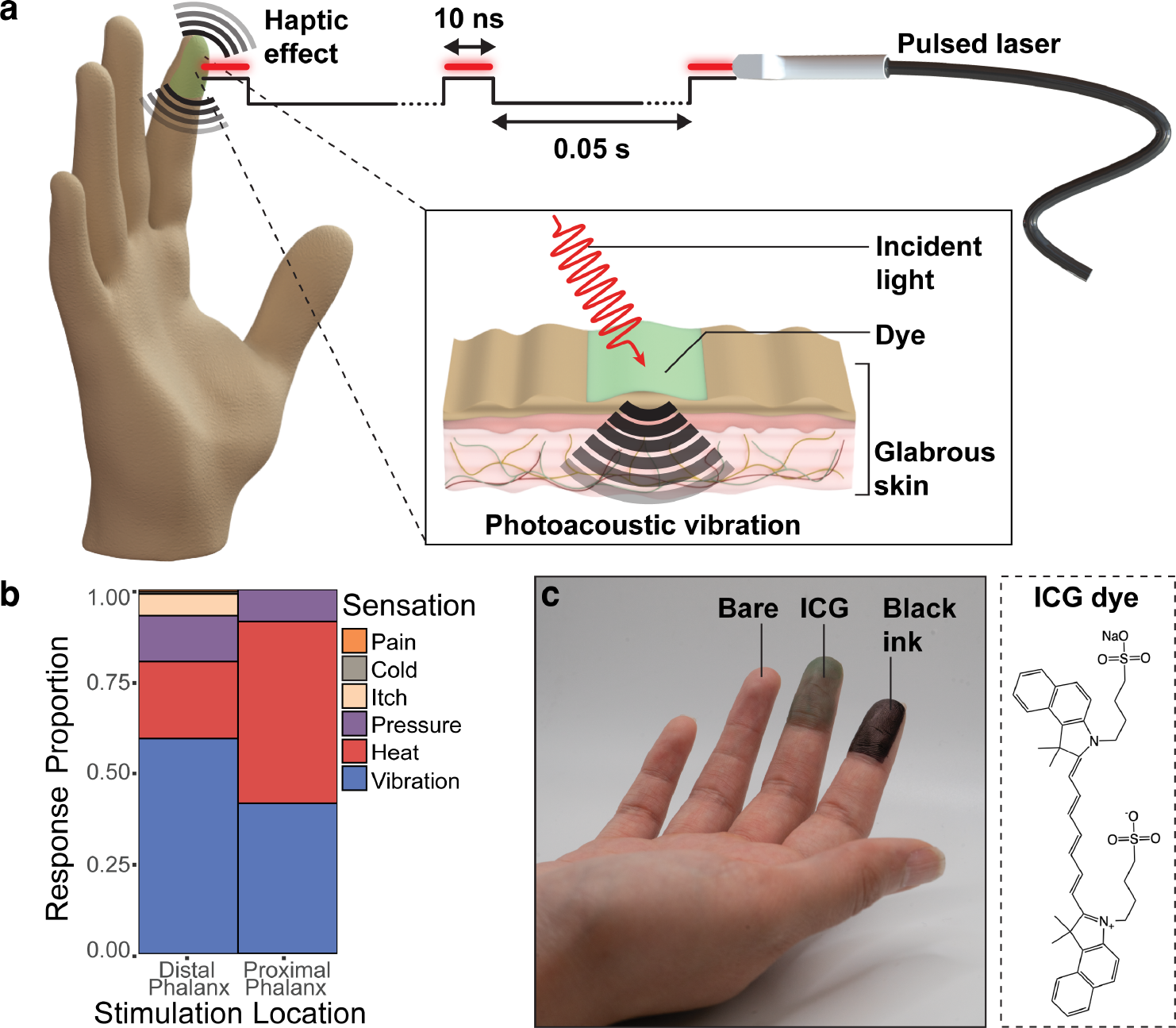
A brief overview of the effect and key result described in this work. **a**, Schematic diagram of the use of pulsed light to generate tactile sensations mediated by the photoacoustic effect. **b**, Summarized proportions of tactile sensations reported by human participants after applying the optical stimulus on the distal (fingertip) and the proximal phalanges. **c**, Photograph of a participant’s fingertips coated with dyes used to sensitize the skin to the photoacoustic effect. The right panel shows the chemical structure of the IR-absorbing dye, indocyanine green (ICG).

The photoacoustic effect generates a wide range of frequencies which depend in part on the width of the optical pulse (tens of nanoseconds in biomedical imaging)[15–17]. These stress waves contain a range of frequencies in the tens of kHz to MHz regime, too high to be detected by afferents of the skin. Nevertheless, the pulses are typically repeated at rates that are well within the range of detection of mechanoreceptors, i.e., 20 Hz to 1 kHz[18]. The duty cycle of the pulses is short enough—often substantially less than 1 percent—that fast dissipation of thermal energy reduces the accumulation of heat in the context of biomedical imaging [19]. Thus, in the current study, along with any future methodology that uses the photoacoustic effect as a means of mechanically activating cells and living tissues, excess thermal accumulation of optical energy is avoided.

We designed a sequence of psychophysical experiments aimed at testing the quality of perception (vibration vs. heat vs. pressure), different dye absorbers (Fig. 1c), spatial effects (direction of movement of the stimulus), and light sources of different powers and wavelengths[20]. Pilot experiments suggested that participants were unable to feel any effect from the pulsed light without the use of a dye. However, when applied to the fingers, a thin film of black ink or translucent indocyanine green (ICG) dye allowed the perception of a strong effect in most participants. For the first battery of experiments (Fig. 2) the light source was a nanosecond optical parametric oscillator (OPO) laser that is commonly used for photoacoustic imaging (Vevo F2 LAZR-X). It delivered 10 ns pulses of 680 or 800 nm wavelength at a repetition rate of 20 Hz (duty cycle = 0.00002%). The beam was spread to illuminate a rectangular area of 2.7 cm^2^ when held 2 cm from a surface. Participants were seated and blindfolded. Audible frequencies emanating from the skin (Supplementary Fig. 1) during the experiments were blocked with foam earplugs combined with noise-canceling headphones.

**Fig. 2.**
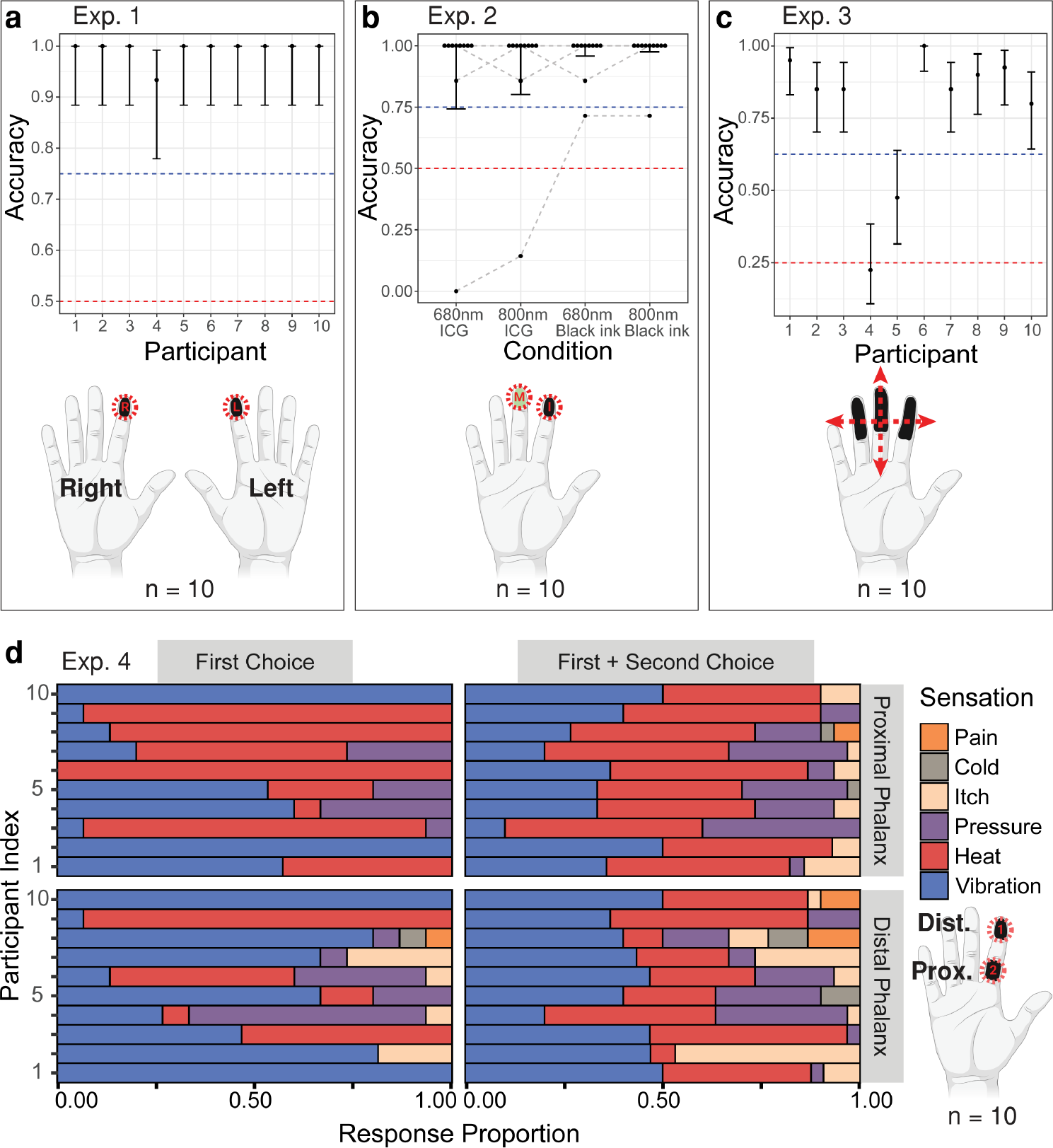
Perception accuracy of the effect in various conditions and effect categorization results produced using OPO system. **a**, Accuracy results of 2AFC experiment to test for detection of stimulus on the index fingers marked with black ink. **b**, Accuracy results for detection of stimulus with different wavelengths and ink types (GLMM estimated marginal means). **c**, Accuracy of 4AFC test for stimulus sweep in four different directions (red arrows) across distal and middle phalanges with black ink. **d**, Proportion of responses for primarily perceived sensations by participants on the proximal and distal phalanges (left panels). Right panels show combination of primary and secondary perceived sensations for proximal and distal phalanges (top and bottom, respectively). Note: All error bars are 95% confidence intervals. The red dotted lines indicate chance performance, and the blue dotted lines depicts the 75% accuracy level typically used in 2AFC or 4AFC experiments to define the detection threshold.

The first experiment was a two-alternative forced choice (2AFC) test in which black ink was applied to the index fingertips of both hands. On each trial, the marked regions of one of the two fingertips was irradiated with 680 nm wavelength stimulus, to test if participants could identify which finger (left or right hand) was stimulated (Supplementary Fig. 2a). Of the ten participants, nine were 100% accurate, and the tenth was accurate on 93% of trials (Fig. 2a). This performance is significantly higher than the 50% that would be expected by chance (separate binomial test for each participant, Bonferroni corrected for multiple comparisons, all *p <* 0.001), and is also significantly higher than 75% accuracy that is usually set by convention to correspond to the detection threshold in 2AFC experiments.

We then sought to investigate whether the stimulus could be perceived with an IR (i.e., nearly invisible) wavelength and a less conspicuous absorber than black ink. We thus turned to ICG, commonly used for tissue staining in biomedical imaging[21, 22], and compared the effect of both absorbers at both 680 and 800 nm (Supplementary Fig. 2b). While overall accuracy was higher with black ink (Wald *Z* = 2.19, *p* = 0.03), this effect appears to have been driven entirely by the results of a single participant (Fig 2b). When the participant’s data are excluded, performance was not significantly influenced by the absorber, wavelength, or their interaction (all *p >* 0.5). Crucially, in all four conditions, participants detected the stimulus more often than would be predicted by chance (*Z* tests on estimated marginal means, all *p <* 0.05; see Supplementary Table 1,2), and in all four conditions all but one participant surpassed the convention of 75% accuracy. It thus appears that the generation of this “phototactile” effect is amenable to a range of absorbers and wavelengths.

Next, we tested the ability of participants to discern the direction of travel of the pulsed beam on the skin. We did so by sweeping it over an inked portion of the dominant hand in one of four directions, and prompting the participants to identify the direction (Supplementary Fig. 2c). Of the ten participants, one was not significantly more accurate than chance (binomial test, *p* = 0.86), one was significantly more accurate than chance (binomial test, Bonferroni-corrected *p* = 0.003) but not more accurate than the typical detection threshold (62.5% for 4AFC), and eight were significantly more accurate than chance (separate binomial test for each participant, Bonferroni corrected for multiple comparisons, all *p <* 0.001) and also more accurate than the typical detection threshold (Fig. 2c). The results from this experiment showed that participants could discern not only the direction of the sweep (i.e. left to right, front to back, etc.), but also the relative position of the laser indicating that the stimulus can be spatially and temporally resolved [23, 24]. Additionally, the experiment demonstrated the ability of participants to detect stimulus across multiple areas of the finger (distal and middle phalanges) and across multiple fingers.

Earlier, we hypothesized that the photoacoustic waves would be perceived as a mechanical stimulus, rather than as, e.g., heat. Indeed, when given a choice of pressure, vibration, itch, pain, heat, or cold, perceived by participants at the distal phalanx (fingertip), vibration was the predominant sensation (Fig. 2d and Supplementary Fig. 2d). Nevertheless, the acoustic waves generated are mediated by transient absorption and thermalization of the optical energy, and thus the possibility remained that the effect could be perceived as thermal. We found that the cumulative temperature increase caused by the pulsed light on the skin (black ink, 680 nm excitation based on the absorption spectrum of black ink (Fig. 3a)) was relatively small, 0.5 ° C, after 10 s of illumination, with larger increases in temperature not achieved until very long dwell times (Fig. 3b). Consistent with that observation, heat did not surpass vibration in terms of the frequency with which it was reported by the participants as the primary sensation, at least at the distal phalanx (primary sensation felt: 59% vibration, 21% heat). However, the sensation was more likely to be perceived as heat when the beam was directed to the skin closer to the palm (proximal phalanx, the primary sensation felt: 42% vibration, 50% heat; *χ*^2^(1)=6.58, p=0.01). This difference in perception may be explained by the reduced concentration of mechanosensory neurons—particularly the Meissner’s corpuscles sensitive to frequencies in the range of the 20 Hz repetition rate of the laser—closer to the palm[25].

**Fig. 3.**
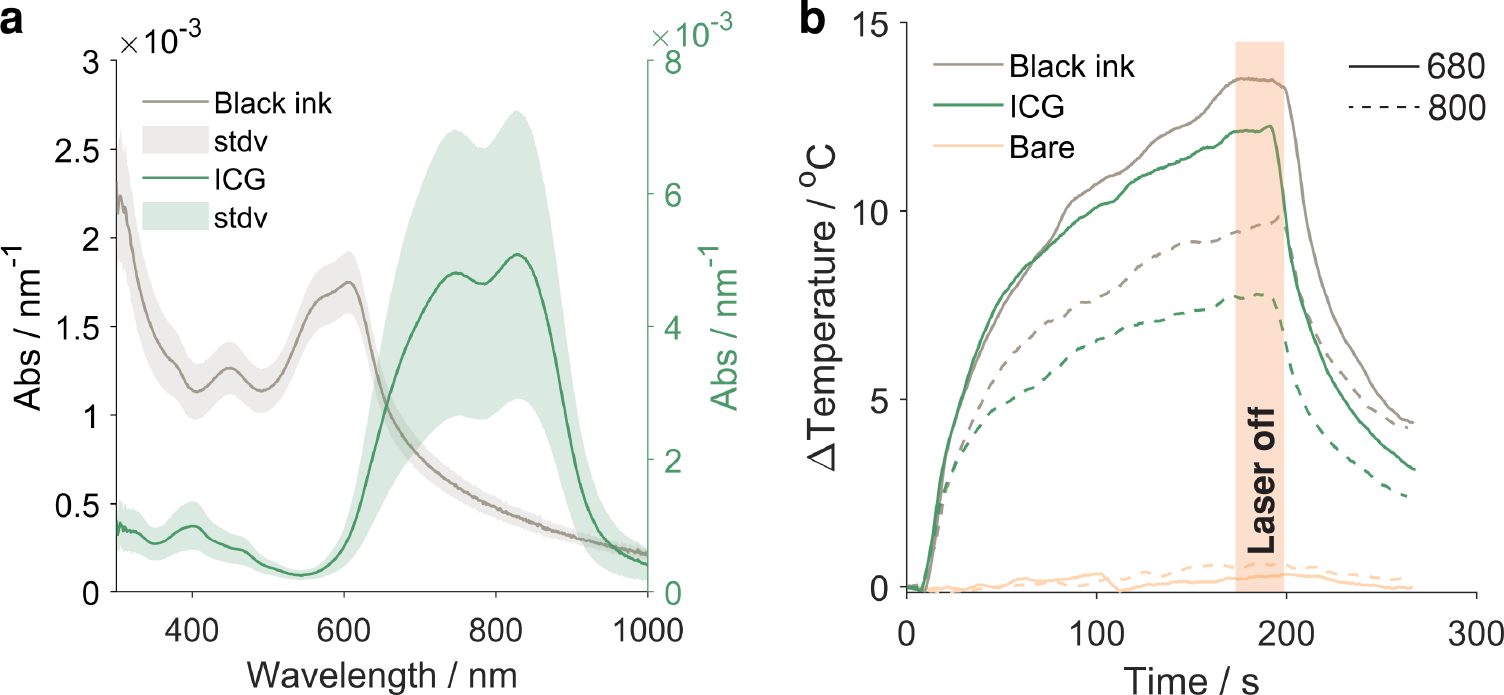
Characterizations of the optical properties of the dyes and the thermal response of the dye-coated skin. **a**, Absorption spectra of black ink and ICG dye spin-cast on glass normalized by film thickness (see **Supplementary Fig. 3**).The absorption spectrum of black ink drawn on glass is shown in **Supplementary Fig. 4. b**, Change in temperature over time with pulsed light of 680 and 800 nm on dyed and bare skin. The laser was powered on for ca. 180 s and then powered off to observe the skin cooling (see **Supplementary Fig. 5 to S7**).

Given the expense (ca. $100,000) and inaccessibility of the medical OPO pulsed laser system, we investigated portable and less expensive light sources. We pulsed these sources (a red laser pointer, 650 nm, and a more powerful miniature diode laser, MDL, 808 nm) at 20 Hz with 6.25 ms pulse width. Using this handheld setup (Fig. 4a and Supplementary Fig. 8), we first conducted a simple 2AFC test on a cohort of 5 participants (Supplementary Fig. 9a). Participants’ accuracy was significantly higher than chance with the MDL (Wald *Z* = 4.39, *p <* 0.001, Supplementary Table 3,4), but not with the red laser pointer (Fig. 4b), probably due to its lower power and smaller spot size. Consistent with the results of our pilot experiments with the OPO laser in which we used no dye (not shown), we found that chopped light from even the MDL laser could not be felt without black ink (2AFC results, Fig. 4c and Supplementary Fig. 9b, Supplementary Table 5,6). Consistent with previous experiments, in the inked condition all but one participant was at or above 75%. Anecdotal evidence from the participants suggests that sensations from the MDL were more likely to be perceived as heat than with the OPO system, possibly owing to a significantly longer duty cycle (i.e., dwell time) due to the homemade mechanical chopper: ca. 12.5% vs 0.00002%.

**Fig. 4.**
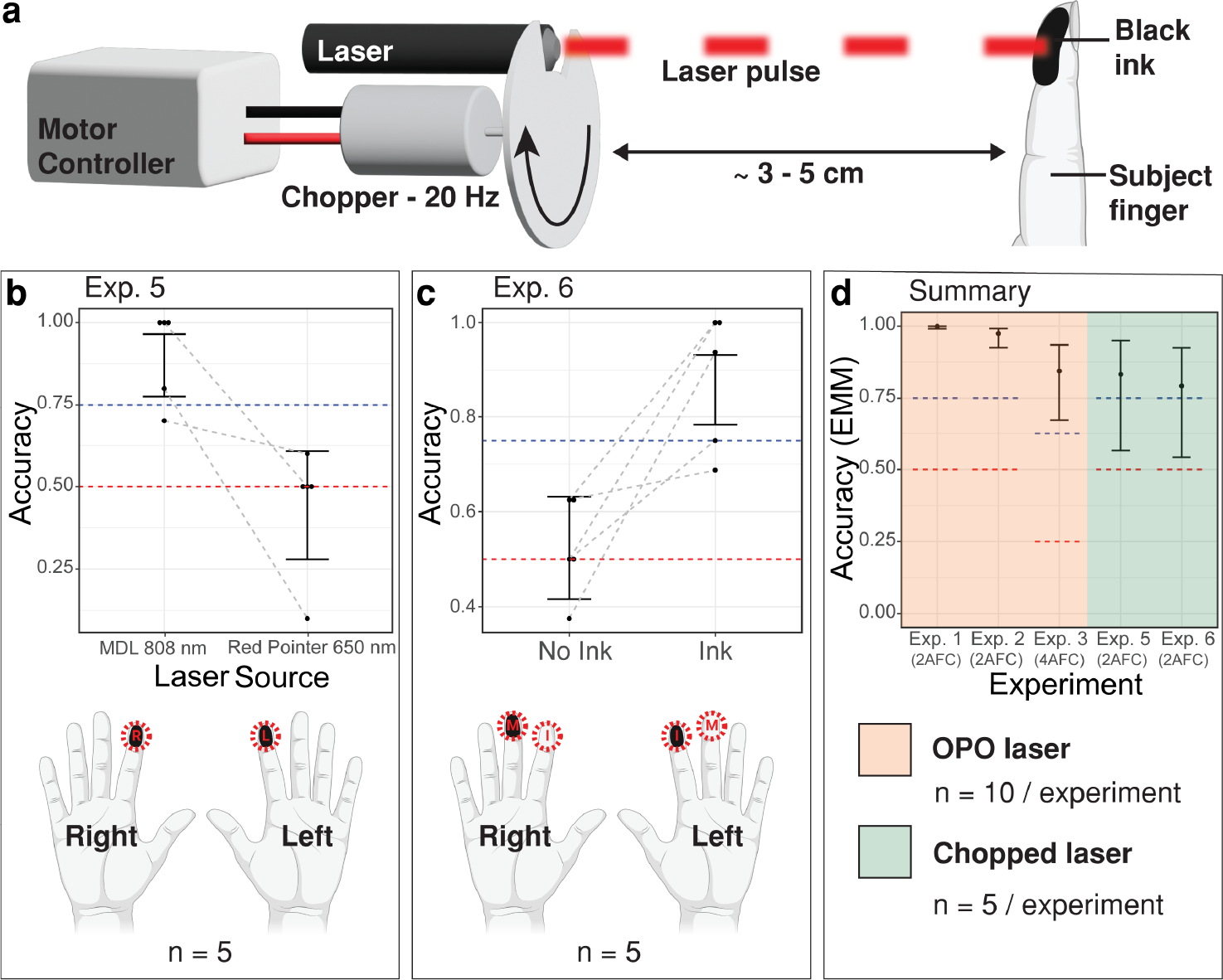
Schematic of the psychophysical experiment and results performed using mechanically chopped light from lower cost laser sources along with an overall experimental summary. **a**, Schematic of the handheld system pulsing light onto an inked finger. **b**, Average participant accuracy to detect stimulus with two different lasers. Error bars are 95% confidence intervals on GLMM estimated marginal means. The red dotted line depicts chance performance (50%), the blue dotted line depicts the accuracy level typically used in 2AFC experiments to define the detection threshold (75%). **c**, Accuracy of participants’ ability to detect the haptic effect with and without ink. Error bar confidence intervals and chance performance are the same as those for **b. d**, Experimental results summary for both OPO and chopped laser systems used in this study. Error bars are 95% confidence intervals on GLMM estimated marginal mean.

The safety concerns of light sources—particularly lasers—are related to potential damage to the retina, burning of the skin, and the generation of reactive oxygen species. PA effects scale with optical power; the radiant exposure of human skin must be below 20 mJ cm^*−*2^ in the near-infrared region. Both of our higher power systems (OPO and MDL-chopper) had divergent beams and the exposure was below this threshold when the distance from the source to the skin surface was 2 cm.

The in situ conversion of optical stimulus into a mechanical sensation, while based on a century-old effect, represents a potentially powerful modality of neural activation in humans. The effect explored here is robust and highly tolerant of a range of physical parameters such as wavelength, pulse rate, pulse width, and dye sensitizer. Key advantages of a system based on in situ generation of mechanical forces in the skin include the ability to project the stimuli without physical contact. Moreover, the use of optical signals opens the door to the simultaneous stimulation of multiple types of afferent fibers. For example, by using overlaid variations in frequency and pulse width, it may be possible to activate multiple types of mechanically and thermally responsive neurons. Advancing such abilities can facilitate the development of “tactile holography” which could have applications in medical training, physical therapy, pain management, complementary healthcare, remote operation, consumer electronics, and various applications of virtual and augmented reality. Moreover, the field of mechanobiology requires tools that can apply forces to cells and tissues in vitro. Such tools generally require physical contact to exert forces on cells and tissues and are often difficult to perform [26, 27]. In particular, in the neurobiology and neuropsychology of touch and interoception, the ability to activate mechanically responsive cells and tissues without genetic engineering (i.e., optogenetics) or physical contact may open the door to new kinds of assays both in vitro and in vivo.

## Notes

### Competing Interest Statement

The authors have declared no competing interest.

