## Supplementary material for "Use of dye sensitizers for increased photoacoustic mechanosensation": Methods and Supplementary Information: SI.pdf

#### Psychophysical experiments

All participants gave informed written consent and all methods were carried out in accordance with the guidelines and regulations approved by the Internal Review Board of UC San Diego. The first four psychophysical experiments (Experiments 1-4) were conducted using a commercial nanosecond optical parametric oscillator (OPO) laser for photoacoustic imaging (Vevo F2 LAZR-X). The applied stimulus had a pulse width of 10 ns with a pulse frequency of 20 Hz. All four experiments used a wavelength of 680 nm, except for Experiment 2, in which 800 nm was also used.

The psychophysical designs for each of the four experiments conducted with the OPO laser are shown in Supplementary Fig. 1. All 10 participants were blindfolded and wearing foam earplugs with noise-cancelling headphones. For each experiment, participants were given the question to be asked prior to the start of the experiment and were prompted to answer the question with a tap on the shoulder at the end of each trial.

**Experiment 1**: The initial experiment design Supplementary Fig. 1a) is a 2-alternative forced-choice test aimed at detecting the stimulus using black ink applied to both index fingers. During each trial, the optical stimulus was presented on one of the index fingers. Participants completed a total of 30 trials, where they were asked, "On which finger did you feel a sensation?" The only acceptable answers were "left index" or "right index". The trials were randomly generated among participants.

**Experiment 2**: An experiment was conducted to investigate the impact of different ink types and wavelengths on stimulus detectability (Supplementary Fig. 1b). Two inks were tested: black ink and indocyanine green (ICG) dye suspended in ethanol (4 mg/mL). The experiment also explored two wavelengths: 680 nm and 800 nm.

For each participant, ICG ink was applied to the distal phalanx of their middle finger, while black ink was applied to the distal phalanx of their index finger, both on their dominant hand. With the two wavelengths, four stimulus conditions were established, representing all possible ink/wavelength combinations: ICG with 680 nm, ICG with 800 nm, black ink with 680 nm, and black ink with 800 nm.

Each participant experienced seven iterations of the four stimulus conditions in a block design, which were randomly ordered, resulting in a total of 28 trials per participant. After each stimulus application, the participants were asked, "On which finger did you feel the sensation?" Acceptable answers were "middle finger" or "index finger."

**Experiment 3**: In this experiment, we employed a 4 alternative forced choice paradigm to examine the identification of motion direction elicited by the applied stimulus. The dominant hand's distal and middle phalanges were marked with black ink. To assess the participant's perception, the investigator swept the stimulus in four distinct directions (Supplementary Fig. 1c): front to back, back to front, left to right, and right to left.

Each participant underwent ten iterations of the four possible directions, presented in a random order using a block design (n = 40 trials per participant). After the stimulus application in each trial, participants were asked, "In which direction did you feel the stimulus sweep?" This procedure allowed us to explore the motion perception patterns effectively.

**Experiment 4**: An experiment was conducted to discern the types of sensations experienced by the participants (Supplementary Fig. 1d). To achieve this, we marked each participant's dominant hand's distal and proximal phalanges with black ink. The stimulus was then randomly applied to either the distal or proximal phalanx in a total of 30 trials. During each trial, the participants were asked "What was the primary sensation you felt?" followed by "What was the secondary sensation you felt?" after the stimulus application. They had six possible sensations to choose from: heat, cool, pressure, vibration, itch, and pain.

Experiments 5 and 6 were conducted using the handheld chopper (Fig. 4). The pulses of light were at a frequency of 20 Hz. There were five participants for experiments 5 and 6.

**Experiment 5**: To determine whether participants could detect stimulation with different lasers (MDL 808 nm 2 W and Red Pointer 650 nm), we conducted an experiment to apply two different laser stimuli on the participants' inked fingers (Supplementary Fig. 9a). The MDL 808 nm laser and Red Pointer 650 nm laser are referred to as laser 1 and laser 2, respectively. The participant's distal phalanges of their index fingers were marked with black ink. There were four possible stimulus conditions: left index with laser 1, right index with laser 1, left index with laser 2, and right index with laser 2. The four conditions were counterbalanced in a total of 20 trials. After each stimulus application, the participants were asked, "On which finger did you feel the sensation?" Acceptable answers were "left index" or "right index."

**Experiment 6**: We conducted an experiment to determine whether participants could detect stimulation without ink. We used the MDL 808 nm laser with the handheld chopper for this experiment. The participant's middle and index fingers on both of their hands were used for the experiment (Supplementary Fig. 9b). Each hand had one of the two fingers marked with black ink (distal phalanx) and the other one bare. The four possible ink placements were counterbalanced across participants, with one participant condition repeated since we had five total participants. The stimulus was applied to each of the four fingers in random order for a total of 8 blocks. After applying the stimulus, the participant was instructed to answer the following question: "On which hand was the sensation felt?"

#### Statistical analysis

**Experiment 1**: We used a separate binomial test for each participant (N = 10) with a 95% confidence interval determined via the Clopper-Pearson method. A Bonferroni correction was applied for multiple comparisons. There were 30 trials per participant (n = 30).

**Experiment 2**: We employed a logistic mixed-effects regression model with the dependent variable being trial outcome (correct/incorrect). The model included fixed effects for ink type (Black ink vs. ICG), wavelength (680nm vs. 800nm), and their interaction, along with a random effect for participant. Each participant (N = 10) experienced seven iterations of the four stimulus conditions in a block design, which were randomly ordered (n = 28). Error estimates were calculated using a 95% confidence interval on the estimated marginal mean accuracy for each condition, which was back-transformed from the logit scale (Supplementary Table 1). Additionally, we conducted a Wald Z statistical test within a generalized linear model framework to assess the significance of differences in detection accuracy attributed to various inks and wavelengths (Supplementary Table 2).

**Experiment 3**: Each participant (N = 10) underwent ten iterations of the four possible directions, presented in a random order using a block design (n = 40) We conducted Bonferroni-corrected binomial tests for each participant, accompanied by a 95% confidence interval.

**Experiment 4**: We computed response proportions separately for primary and secondary sensations within each sensation type. Subsequently, we combined the proportions of primary and secondary responses and calculated overall response proportions for each sensation type encompassing both primary and secondary sensations. To test whether heat was experienced more often at the proximal than the distal phalanx, we combined the proportions of all sensations other than heat and ran a 2-by-2 chi-squared test of independence (N = 10, n = 30).

**Experiment 5**: We fit a logistic mixed effects regression with dependent variable of trial outcome (correct/incorrect), fixed effect of laser (MDL 808 nm vs. Red Pointer 650 nm), and random effect of participant. Participants (N = 5) experienced each stimulus condition 5 times (n = 20). Error was calculated with a 95% confidence interval on estimated marginal mean of accuracy per condition, back-transformed from the logit scale. Additionally, we conducted a Wald Z test of the laser term to assess the significance of differences in detection accuracy attributed to the different lasers.

**Experiment 6**: We fit a logistic mixed effects regression with dependent variable of trial outcome (correct/incorrect), fixed effect of ink (present/absent), and random effect of participant. Each of the four fingers were stimulated in random order, with 8 blocks total per participant (N = 5, n = 32). Error was calculated with a 95% confidence interval on estimated marginal mean of accuracy per condition, back-transformed from the logit scale. Additionally, we conducted a Wald Z test of the ink term to assess the significance of differences in detection accuracy attributed to the presence of ink.

#### Experimental setup

**Experiments 1-4:**A chair and bench top were prepared for the human subject, fitted with a platform to place their hands on. The optical parametric oscillator laser of a VisualSonics Vevo F2 LAZR-X was connected to the medium size fiberoptic cable. The two ends of the cable were taped together for easier handling.

The subjects were asked to place in foam earplugs and a blindfold. The subjects' hands were cleaned from any residues and the dye was applied corresponding to each experiment. Two coats of ICG Dyes were applied using a cotton swab and allowed to dry between coats. The black ink was directly applied onto the subjects' finger/s using a permanent marker (Sharpie Fine Point - Black). Noise-canceling headphones (RUNOLIM WH301A) were placed on the subjects' ears. A Python script was used to randomize the stimulation parameters.

**Experiments 5-6:** The subject preparation for experiments 5-6 was identical to that of experiments 1-4. The laser chopper setup was built using a simple DC motor controlled using an Arduino microcontroller fitted with a motor control shield. Parts were designed in a 3D CAD software and printed on an FDM 3D printer using poly-lactic acid (PLA) and flexible thermoplastic polyurethane (TPU) filaments. The laser socket was made to fit the various laser sizes examined allowing for easy swapping during the experiment. For thin fiber optic cables custom adapters were made to ensure a tight fit and minimize deviation.

#### Characterization

I Dye absorbance:

ICG solid was dissolved into ethanol (4 mg/ml), and filtered through a 0.45 $\mu$m, 13 mm filter to remove aggregates. For a controlled comparison, black ink was extracted from a permanent marker (Sharpie Fine Point - Black) by placing the tip of a fresh marker into a vial of ethanol (20 ml) overnight. Glass slides (1" x 1") were cleaned in a sonication bath using the following wash sequence (Alconox solution, water, isopropanol). The slides were then dried and treated in a plasma cleaner (400 mTorr air, 3 minutes, 30 W) to activate the surface. The solution was spun on the glass slides at 600 rmp and 300 rpm/s. Three samples were prepared of each dye.

The absorption spectra of each sample were obtained between 300 nm and 1000 nm (Aligent Cary UV-Vis Spectrophotometer). The thickness of the samples was evaluated at two points using ellipsometery (J.A. Woollam M-2000D Spectroscopic Ellipsometer). The thickness was used to normalize the absorption spectra. The absorption spectrum of a permanent marker drawn on a glass slide was also obtained to rule out any inadvertent changes to the marker ink during extraction.

II Thermography:

A willing subject's index and ring fingers were coated with black ink and ICG respectively, the middle finger was set as the control (bare). A thermographic camera (HIKMICRO Pocket 2) was affixed on a stand to measure the temperature of the finger tip at the laser's area of incidence. The ambient temperature was input into the camera settings and the emissivity setting was obtained for each finger by validating the finger with a thermocouple (Fluke t3000 FC). Each finger was illuminated with the laser source using the OPO with identical wavelengths and pulse settings used in the psychophysical experiments for ca. 3 minutes. The laser was then turned off to observe the cooling effect. The thermographic data was extracted from the video footage using MATLAB Image Processing Toolbox and Computer Vision Toolbox. The data was smoothed using a moving mean (20 samples) and plotted against the time signature of the frame. Selected frames were extracted at a chosen time stamps for visualization of the process in Supplementary Fig. 5-7.

III Audio measurements:
A willing subject's index, middle, and ring finger were prepared in a manner similar to the thermography experiments. The hand was placed in a box padded with noise insulation. A lapel microphone without a pop filter was placed inside the box behind the location of the laser to avoid any photoacoustic effects on the microphone surface. The audio jack of the microphone was connected to a SONY Alpha 7R camera. The laser was applied on the different dyes and bare finger while the audio was recording. This experiment was repeated for bare and inked (ICG and black ink) fingers for both wavelengths. To capture the ambient noise generated by the laser, the laser probe was pointed away from the subject and held far away ($> 1$ m) from any surfaces. The audio was collected from the video recordings. A 10kHz high pass filter (24 dB) was applied to the audio data to remove the background machine noise.

IV Chopped laser power measurements:
The laser chopper was affixed on a vertical stand and ensured it was level. A digital power meter (Thorlabs PM100D) was connected to a compatible photodiode power sensor (Thorlabs S120VC). The photodiode was placed below the laser and in its direct line of sight. The sensor was set to the wavelength appropriate to the laser to be tested. The distance between the sensor and source was evaluated with calipers and varied incrementally at 10 mm per increment. The measurements was taken with the chopper turned on, and when it was turned off. This experiment was repeated for the laser pointer and the MDL 808 nm source.

V OPO laser power measurements:
An oscilliscope (TDS2022C) was connected to a biased photodetector (Thorlabs DET10A2 200 - 1100 nm). The OPO laser was pointed directly at the center of the detector sensor and was set to pulse 680 nm wavelength light. The oscilloscope captured the sensor power readings (Supplementary Fig. 10). Confirmed by the manufacturer, the OPO laser, while at a 20 Hz pulse repetition rate, skips a pulse every 4 pulses, resulting in 16 pulses every second.

**Supplementary information.** All supplementary information is provided in an appended document.

**Acknowledgments.** This project was supported by the National Science Foundation BRITE-Pivot program under award number CMMI-2135428 to D.J.L. Additionally, L.L.B. acknowledges the support provided by the National Science Foundation Graduate Research Fellowship Program under Grant DGE-2038238 and by the Achievement Rewards for College Scientists (ARCS) Foundation. The authors acknowledge equipment support via NIH S10 OD032268. The hand schematics in Figures 2 and 4 and Supplementary Figures 2 and 9 were created using BioRender.com.

### Declarations

- Competing interests
  The authors declare no competing interests.
- Ethics approval
  This study was approved by the Internal Review Board at UC San Diego.
- Consent to participate
  Participants provided written informed consent to participate in this study.
- Availability of data and materials
  The data that support the findings of this study are available from the corresponding author upon reasonable request.
- Code availability
  The code used to generate random trials and to analyze thermographic video footage in this study are available from the corresponding author upon reasonable request.
- Authors' contributions
  - Conceptualization: DJL
  - Data curation: LLB, TR, NR
  - Methodology: LLB, TR, NR
  - Investigation: LLB, TR, BQ, DJL, LF, LS, GE, YQ
  - Visualization: LLB, TR, NR
  - Funding acquisition: DJL, JVJ
  - Project administration: DJL, JVJ
  - Supervision: DL, JVJ, RR
  - Writing – original draft: DL, LLB, TR
  - Writing – review & editing: LLB, TR, NR, JVJ, DJL

### Supplementary Information

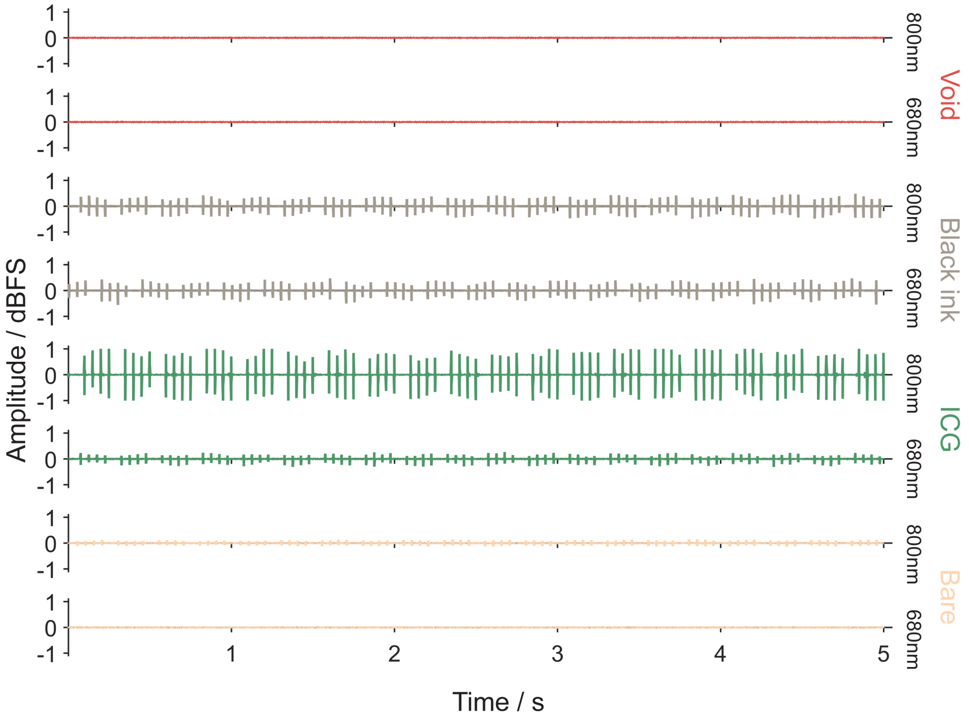


**Supplementary Fig. 1: Graphical representation of the audio signals generated by OPO pulses (800 nm and 680 nm) incident on inked (black ink and ICG) fingers, bare finger, and void.** Note: The OPO laser skips a pulse every 4 pulses (16 pulses per second), though still pulsing at 20 Hz (see Supplementary Fig. 10).

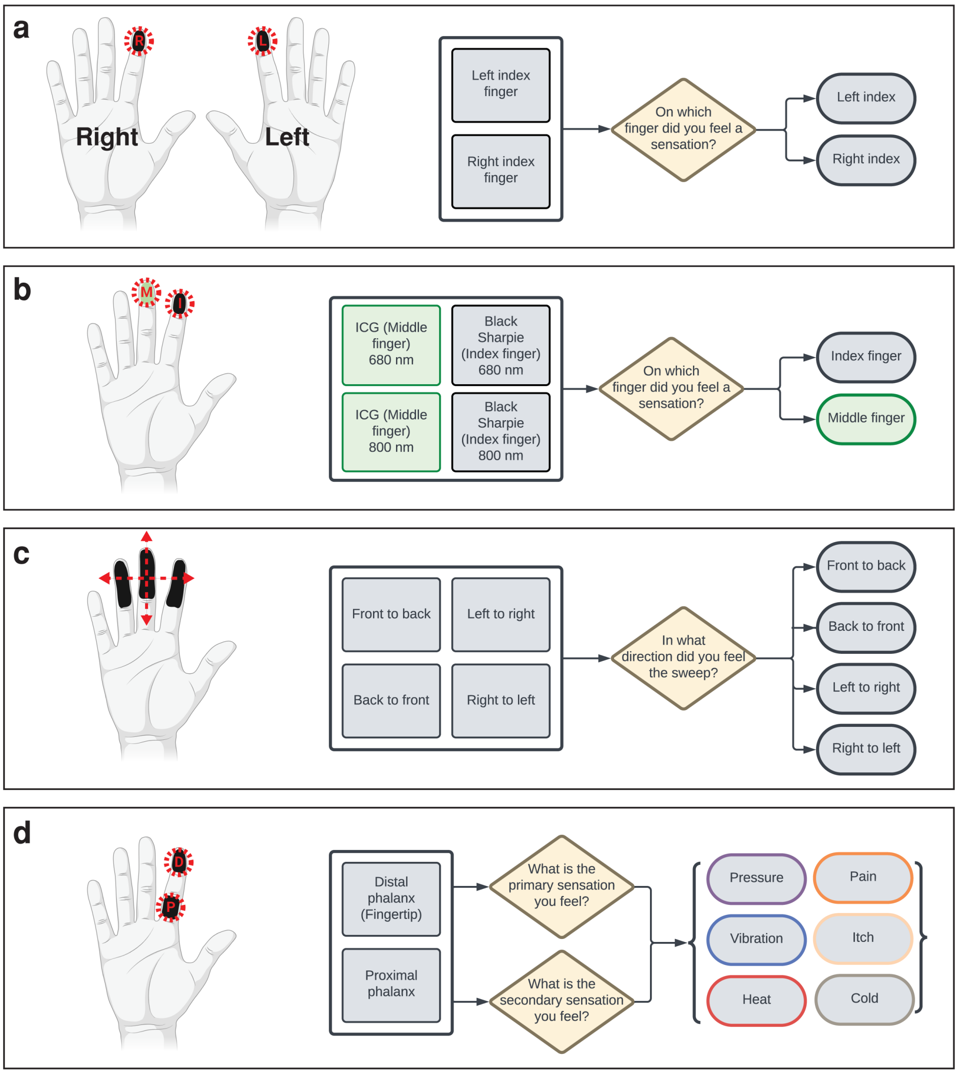


**Supplementary Fig. 2 Schematic (left) and flowchart (right) description of OPO laser experiments for: a,** Experiment 1 **b,** Experiment 2 **c,** Experiment 3 **d,** Experiment 4.

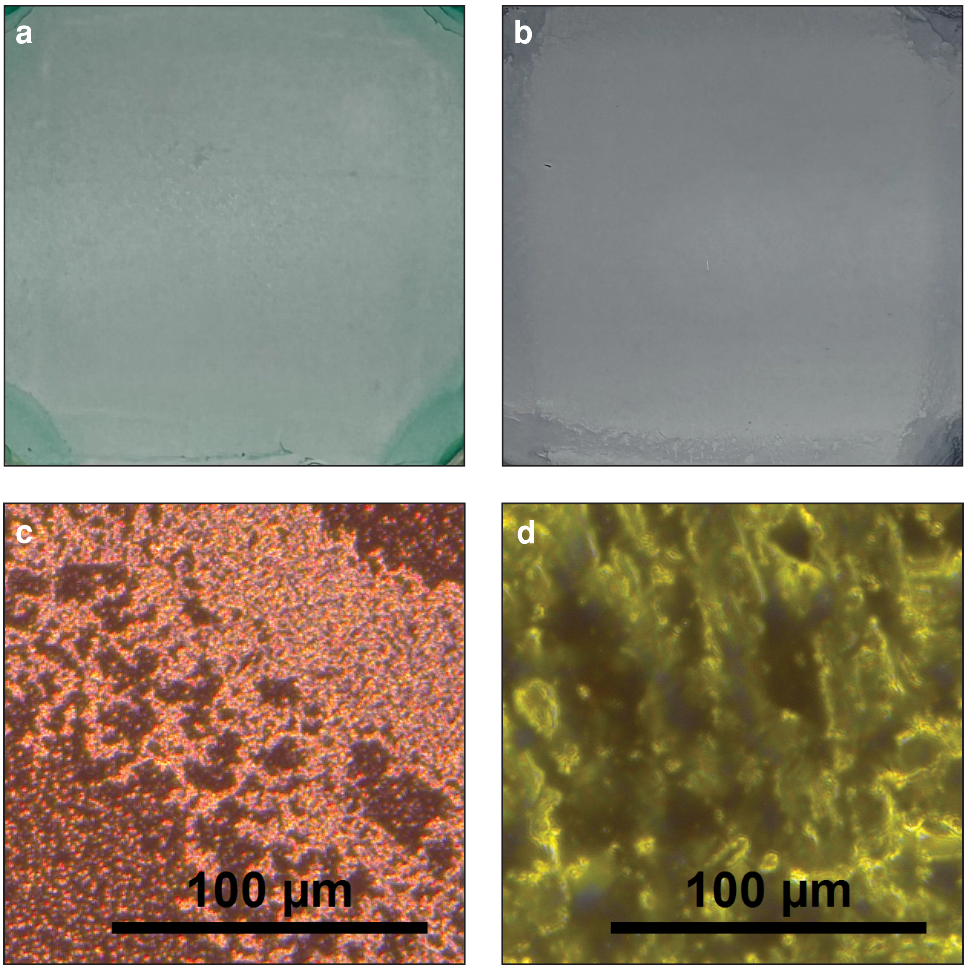


**Supplementary Fig. 3 Photographic and optical microscopy images of spin-cast dye films. a,** Photograph of ICG film on glass substrate. **b,** Photograph of black ink film on glass substrate. **c,** Corresponding ICG film dark field optical microscopy image. **d,** Corresponding black ink film dark field optical microscopy image.

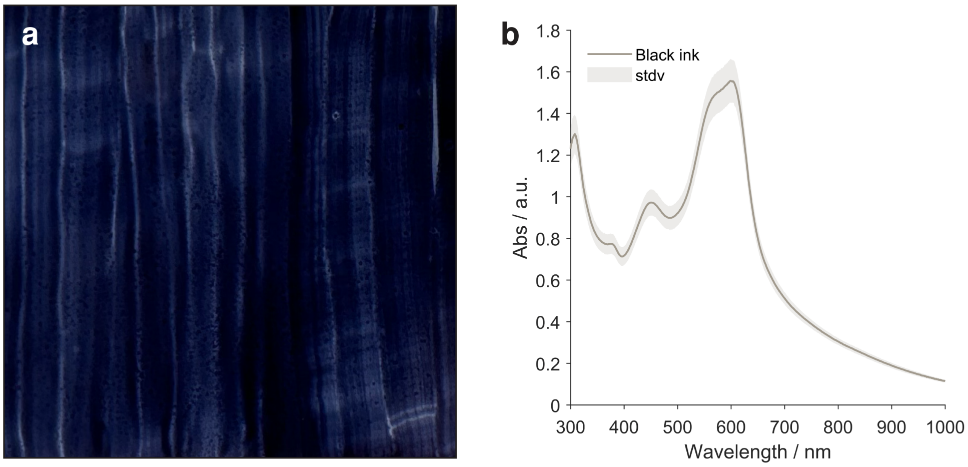


**Supplementary Fig. 4 Photographic and absorbance spectrum representations of black ink drawn on a glass slide. a,** Photograph of black ink film hand drawn on glass substrate. **b,** Absorbance spectrum of black ink drawn film not normalized by thickness.

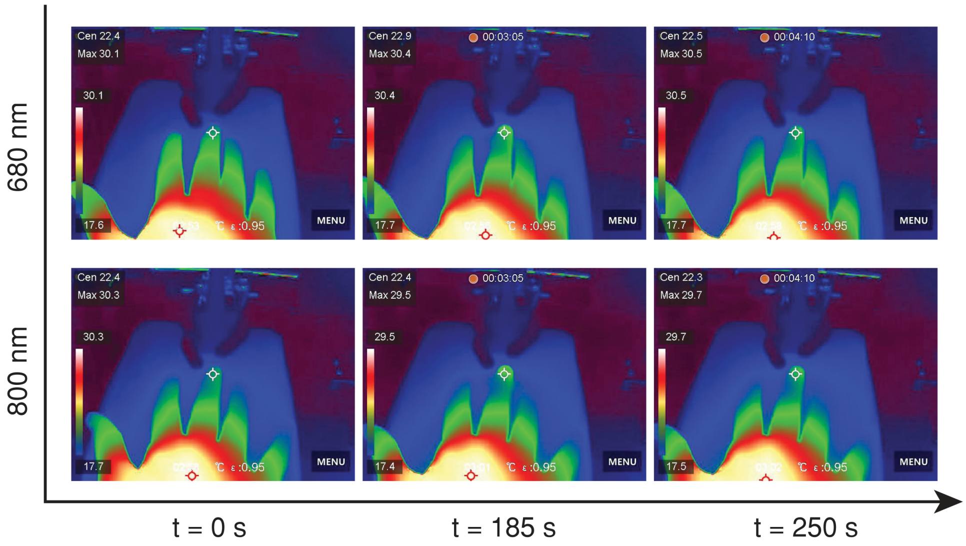


**Supplementary Fig. 5** **Thermographic images during laser stimulation on bare skin.**

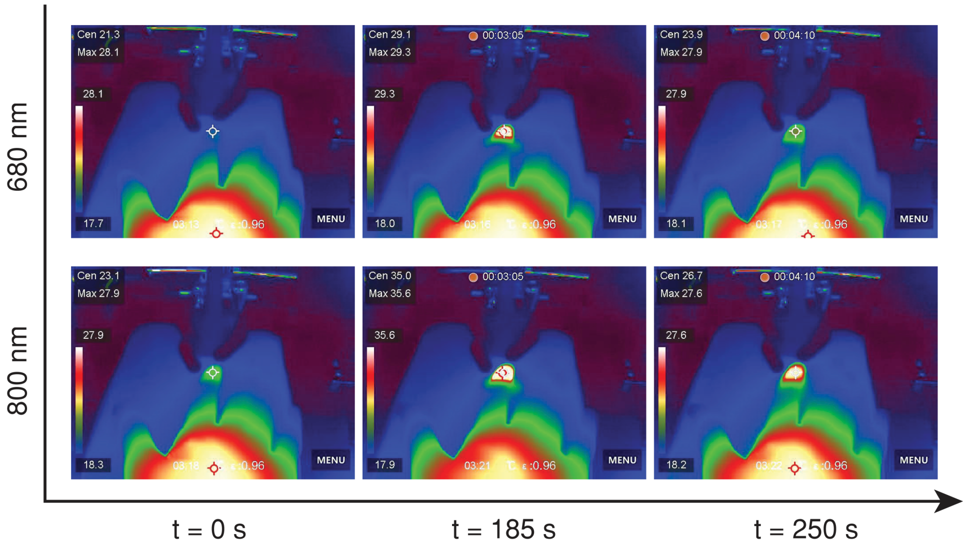


**Supplementary Fig. 6** **Thermographic images during laser stimulation on skin coated with ICG.**

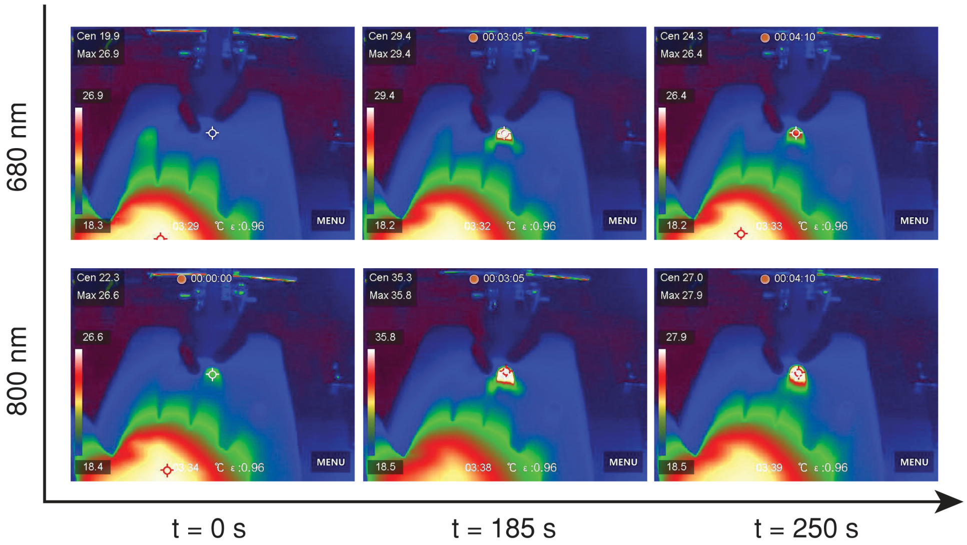


**Supplementary Fig. 7  Thermographic images during laser stimulation on skin coated with black ink.**

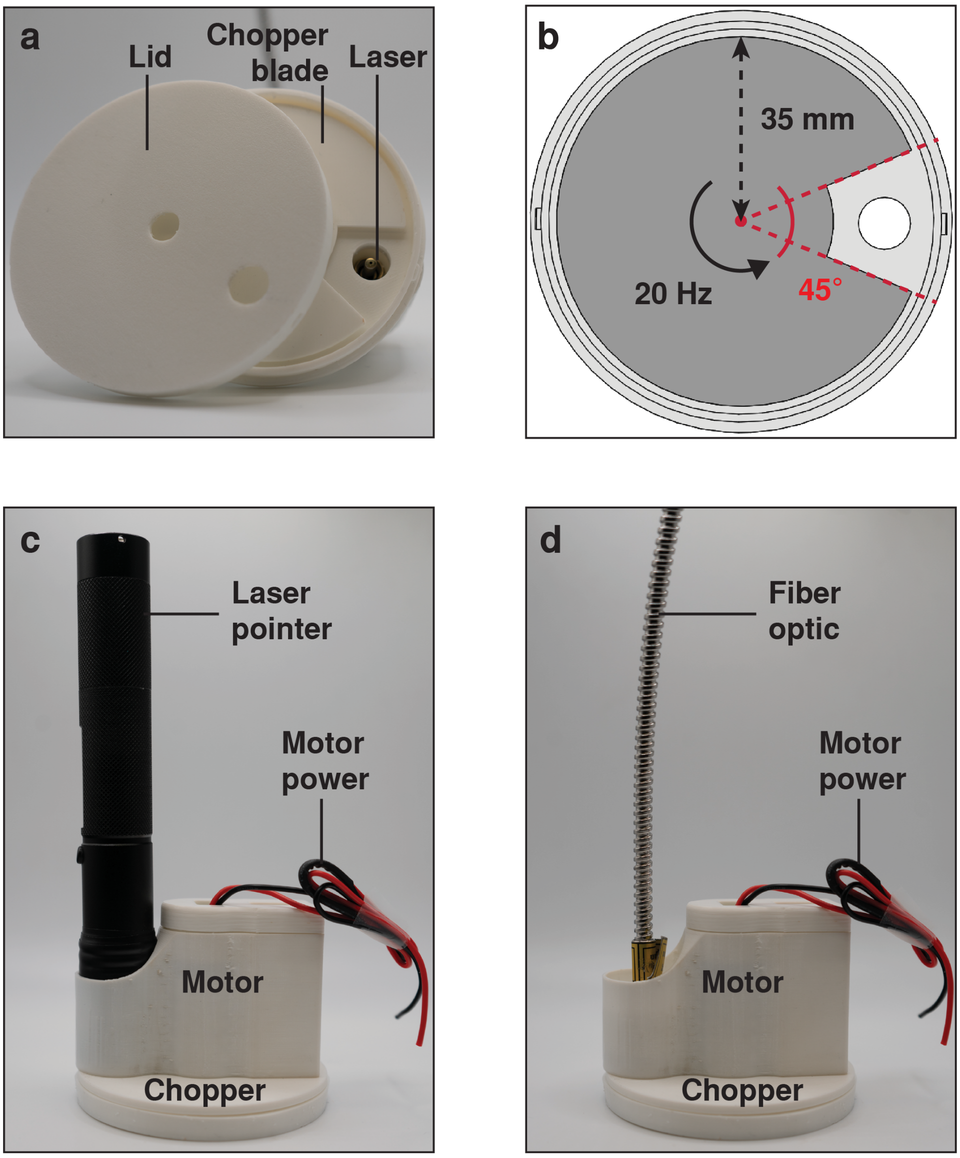


**Supplementary Fig. 8 Photographic and schematic details of the hand-held laser chopper.** **(A)** Chopper assembly showing the laser aperture, chopper blade, and protective lid. **(B)** Chopper blade schematic showing the angle of the chopper window (aligned with aperture) and the diameter of the blade. **(C)** Handheld chopper assembled with laser pointer in the socket. **(D)** Handheld chopper assembled with fiber optic cable in the socket.

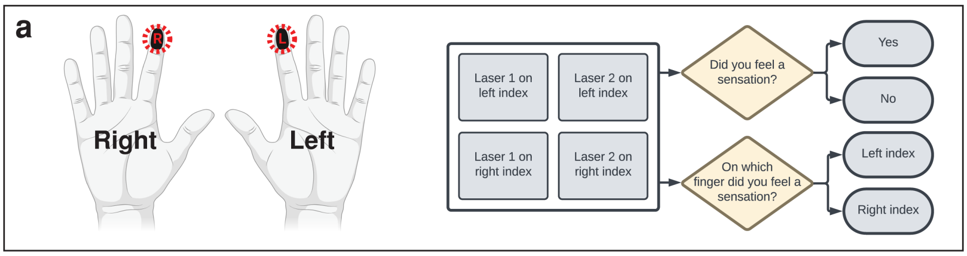


**Supplementary Fig. 9 Schematic (left) and flowchart (right) description of handheld laser experiments for: (A)** Experiment 5 **(B)** Experiment 6.

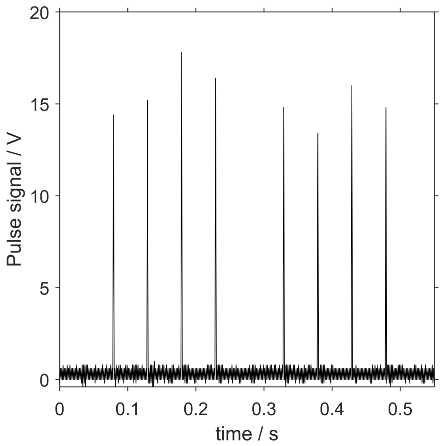


**Supplementary Fig. 10 OPO laser power pulse signal.**

**Table 1 Exp. 2: General linearized mixed model (GLMM) results.**

| Term | Coefficient | Std. Err. | Wald $Z$ | $p$ value |
|---|---|---|---|---|
| Intercept | 7.37 | 3.22 | 2.29 | 0.02 |
| Wavelength (800 nm) | 0.47 | 0.98 | 0.48 | 0.93 |
| Ink (Black) | 2.252 | 1.15 | 2.19 | 0.03 |
| Ink * Wavelength | 0.15 | 1.49 | 0.10 | 0.92 |

**Table 2 Exp. 2: GLMM Estimated Marginal Means (EMMs) of each condition.** Values are back-transformed from the logit scale. Tests are performed on the logit scale

| Wavelength | Ink | Accuracy EMM | Accuracy EMM 95%CI | Wald $Z$ | $p$ value |
|---|---|---|---|---|---|
| 680 | ICG | >0.99 | [0.74, 1.00] | 2.29 | 0.02 |
| 800 | ICG | >0.99 | [0.80, 1.00] | 2.38 | 0.02 |
| 680 | Black ink | >0.99 | [0.96, 1.00] | 2.86 | 0.004 |
| 800 | Black ink | >0.99 | [0.98, 1.00] | 3.01 | 0.003 |

**Table 3 Exp. 5: General linearized mixed model (GLMM) results.**

| Term | Coefficient | Std. Err. | Wald $Z$ | $p$ value |
|---|---|---|---|---|
| Intercept | 2.28 | 0.53 | 4.30 | <0.001 |
| Laser Pointer | -2.53 | 0.58 | 4.39 | <0.001 |

**Table 4 Exp. 5: GLMM Estimated Marginal Means (EMMs) of each condition.** Values are back-transformed from the logit scale. Tests are performed on the logit scale.

| Laser | Accuracy EMM | Accuracy EMM 95%CI | Wald $Z$ | $p$ value |
|---|---|---|---|---|
| MDL 808 nm | 0.91 | [0.78, 0.96] | 4.30 | $< 0.001$ |
| Laser pointer | 0.44 | [0.28, 0.61] | 0.72 | 0.47 |

**Table 5 Exp. 6: General linearized mixed model (GLMM) results.**

| Term | Coefficient | Std. Err. | Wald $Z$ | $p$ value |
|---|---|---|---|---|
| Intercept | 0.10 | 0.22 | 0.45 | 0.66 |
| Ink | 1.85 | 0.41 | 4.55 | <0.001 |

**Table 6 Exp. 6: GLMM Estimated Marginal Means (EMMs) of each condition.** Values are back-transformed from the logit scale. Tests are performed on the logit scale.

| Laser | Accuracy EMM | Accuracy EMM 95%CI | Wald $Z$ | $p$ value |
|---|---|---|---|---|
| No ink | 0.53 | [0.42, 0.63] | 0.45 | 0.65 |
| Ink | 0.88 | [0.78, 0.93] | 5.76 | <0.001 |

**Table 7 Dye film thicknesses obtained by ellipsometry.** Each film sample was measured at two separate points denoted by the header: "thickness".

| | Thickness 1 (nm) | Thickness 2 (nm) | Mean thickness (nm) |
|---|---|---|---|
| | | ICG | |
| Sample 1 | 76.52 | 60.09 | 68.31 |
| Sample 2 | 65.79 | 71.03 | 68.41 |
| Sample 3 | 48.44 | 45.07 | 46.76 |
| | | Black ink | |
| Sample 1 | 86.05 | 68.63 | 77.34 |
| Sample 2 | 68.26 | 57.26 | 62.76 |
| Sample 3 | 65.37 | 68.37 | 66.87 |

**Table 8 Power measurement of chopped and continuous laser pointer beam over a range of distances (with beam diameter of 3.25mm – distance invariant).**

| Distance (mm) | Laser power (mW) | Chopped laser power (mW) |
|---|---|---|
| 46 | 87.6 | 11.8 |
| 56 | 86.7 | 11.7 |
| 66 | 87.1 | 11.3 |
| 76 | 87.5 | 11.3 |
| 86 | 87.3 | 11.5 |
| 96 | 87.6 | 11.2 |
| 106 | 86.7 | 10.9 |
| 116 | 87.1 | 10.7 |

**Table 9 Power measurement and diameter of MDL 808 nm beam over a range of distances.** Power reported measured for chopped and continuous beams.

| Distance (mm) | Diameter (mm) | Laser power (mW) | Chopped laser power (mW) |
|---|---|---|---|
| 40 | 8.1 | HI >> 130 | 35.1 |
| 50 | 12.5 | HI | 30.1 |
| 60 | 15.3 | HI | 27.5 |
| 70 | 17.9 | HI | 25.9 |
| 80 | 19.1 | HI | 24.5 |
| 90 | 22.1 | HI | 23.9 |
| 100 | 25.0 | HI | 22.2 |
| 110 | 25.9 | HI | 20.5 |

**Table 10 Total rectangular area of OPO laser beam projected over different distances.**

| Distance (mm) | Area ($mm^2$) |
|---|---|
| 20 | ca. 270 |
| 100 | ca. 990 |
